# Network pharmacological analysis for the identification of the molecular mechanism of *Lilium brownii* (Baihe) against gastric cancer: 3-Demethylcolchicine targeting androgen receptor

**DOI:** 10.1101/2023.04.08.536129

**Authors:** Zi-Yi An, Wen-Hao Zhang, Xiao-Gang Hu, Le-Qi Yuan, Wei-Lin Jin

## Abstract

*Lilium brownii* (Baihe) contains several bioactive compounds with anti-cancer properties. This study aimed to predict the anticancer targets and related pathways of Baihe for the treatment of gastric cancer (GC) by using network pharmacology and to further explore its potential mechanism in GC. The active compounds and their target proteins were screened from the Traditional Chinese Medicine Systems Pharmacology Database and Analysis Platform (TCMSP). The OMIM, CTD, and GeneCards databases provided information on GC-related targets. After the overlap, the targets of Baihe against GC were collected. The STRING network platform and Cytoscape software were used for protein–protein interaction (PPI) network and core target investigations. Network pharmacology predicted that the principal targets were retrieved from the Starbase database in connection with the GC overall survival. Molecular docking was also used to validate Baihe and the targets’ high affinity. Finally, the DAVID online tool was used for the overlapping target Gene Ontology (GO) and Kyoto Encyclopedia of Genes and Genomes (KEGG) pathway enrichment analyses. The TCMSP database showed that Baihe has seven bioactive components. Apoptosis and p53 signaling pathways were primarily enriched in overlapping genes according to KEGG analysis. Androgen receptor (AR) was identified as a major target by combining the PPI network, KEEG enrichment, and target gene prognostic analysis. Molecular docking results verified that the Baihe’s 3-demethylcolchicine has a high affinity for the GC target AR. Based on the results of network pharmacology analysis based on data mining and molecular docking methods, the multi-target drug Baihe may be a promising therapeutic candidate for GC, but further in vivo/ex vivo research is required.

**Graphical Abstract:** 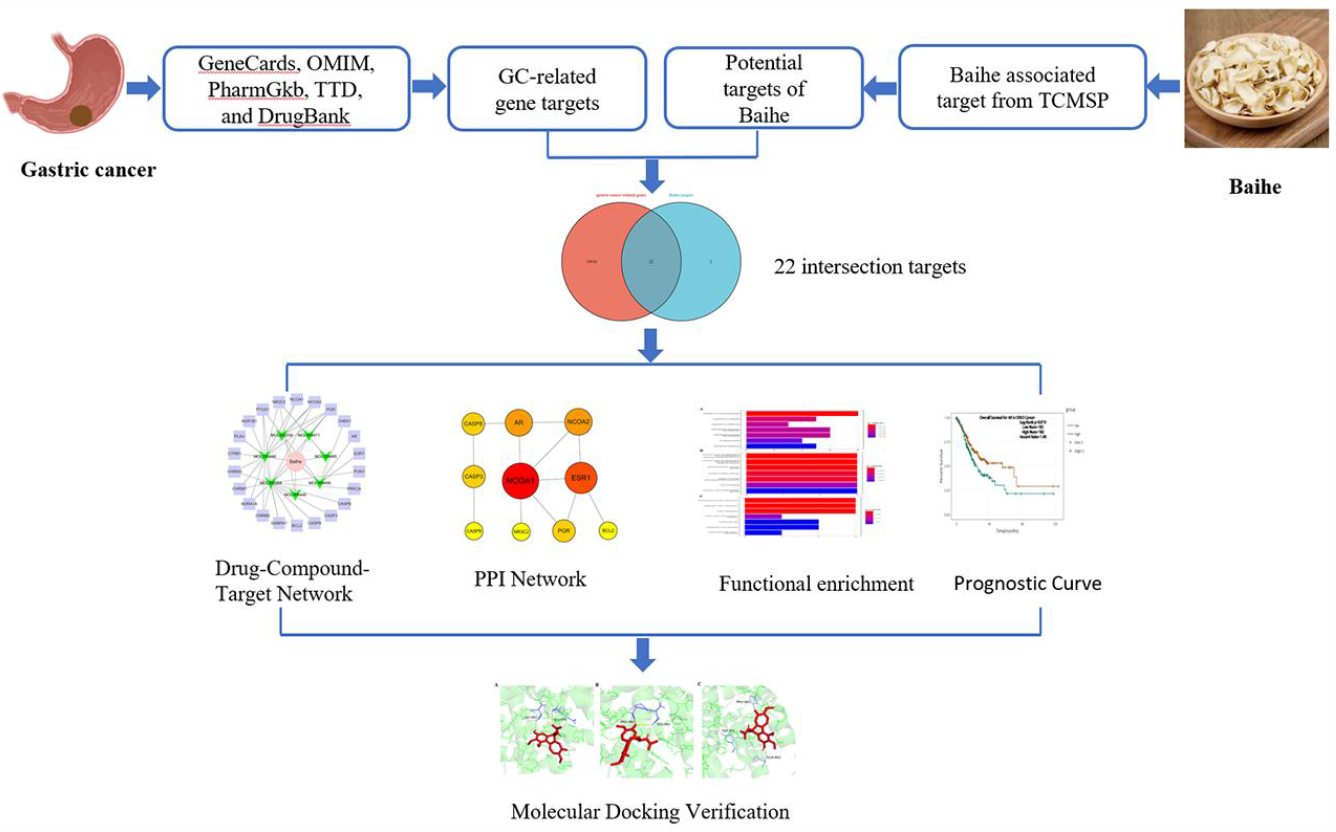

## 1. Introduction

Gastric cancer (GC) is the fifth most common malignancy and the third leading cause of cancer-related death worldwide^1^. The mortality rate for GC has remarkably decreased because of medical advancements. Unfortunately, the prognosis for patients with advanced GC remains poor, and only a small percentage of patients may benefit from current therapy. Additionally, many patients struggle to cope with the side effects of chemotherapy and radiotherapy, and even targeted therapy or tumor immunotherapy may result in sickness among these patients^2^. Thus, novel, safe, and efficient treatments need to be developed for GC.

*Lilium brownii* is a herbaceous perennial bulb plant of the Liliaceae family, and it is widely cultivated in China. This plant was initially mentioned in the Shen Nong Ben Cao (神农百草) and is now formally included in the Chinese Pharmacopoeia. The bulb, also known as “Baihe,” has long been used as an herbal treatment for chronic gastritis, whooping cough, pneumonia, bronchitis, and cough. Based on the ancient Chinese medical system, Baihe may nourish yin and moisten the lungs, eliminate heat, and purify the body, and regulate the spleen and stomach. They are also a significant components of our nutrition and are frequently utilized as functional food^3^. Baihe contains several bioactive compounds with anti-tumor properties^4-6^. For instance, pure lily polysaccharides increase macrophage phagocytosis, splenocyte proliferation, and cytokine (TNF-, IL-2, IL-6, and IL-12) production while dramatically inhibiting Lewis’s lung cancer development and transplanted melanoma^7^. Limited information is available about the bioactive components in Baihe that may have anti-gastric cancer benefits, possible targets, and molecular processes.

Traditional Chinese medicine (TCM) theory is comprehensive and organized. Based on TCM theory, various compounds interact with various cellular targets via multiple pathways to provide therapeutic effects^8, 9^. Network pharmacology is a methodical approach that integrates computer science, bioinformatics, and molecular biology into the pharmacological analysis of TCM to investigate its mechanism of action. It also serves as a theoretical foundation for future studies on natural medicines^1, 10^.

In the present study, an active ingredient-target-disease network was constructed to screen the active components of Baihe and their targets in GC, predict the relevant targets and signaling pathways, and investigate the therapeutic mechanism of Baihe in GC.

## 2. Methods

### 2.1 Construction of drug–compound–target network

The active ingredients in Baihe were screened by integrating oral bioavailability (OB ≥ 30%) and drug-likeness (DL ≥ 0.18), and their targets were screened out in the Traditional Chinese Medicine Systems Pharmacology (TCMSP) Database (https://old.tcmsp-e.com/tcmsp.php). The targeted genes in gastric cancer were obtained from GeneCards (https://www.genecards.org/), OMIM (https://www.omim.org/), PharmGkb (https://www.pharmgkb.org/), TTD (https://db.idrblab.net/ttd/), and DrugBank (https://go.drugbank.com/) databases. The intersected targeted genes were obtained by cross-matching. The drug–compound– target network was visualized using Cytoscape (version 3.5.1).

### 2.2 Construction of the target protein–protein interaction (PPI) network

The interaction between the target proteins was determined by putting the intersection genes into the STRING database (https://cn.string-db.org/) to construct PPI networks. The nodes in the network meet the minimum required interaction score of 0.900, and disconnected nodes were hidden. The downloaded data were visualized using Cytoscape software for better understanding.

### 2.3 Functional enrichment and Kaplan-Meier survival analysis

The pathways that the target proteins may be involved in and their effects on the prognosis of patients with gastric cancer were determined by conducting Kyoto Encyclopedia of Genes and Genomes (KEGG) and Gene Ontology (GO) analysis, and the results were visualized using the R “clusterProfiler” package on Hiplot (https://hiplot.com.cn/home/index.html). Survival analysis was carried out in gastric cancer, and the Kaplan–Meier curves of target genes were obtained based on the Starbase database. (https://starbase.sysu.edu.cn/index.php). Statistical significance was considered at p value < 0.05.

### 2.4 Molecular docking and visualization of key protein and its ligand

The mode of binding of small molecule ligands to large molecule proteins was demonstrated by conducting molecular docking and visualization. The structure of AR protein was obtained in the PDB database (https://www1.rcsb.org/), and the structure of 3-demethylcolchicine is detailed in the PubChem database (https://pubchem.ncbi.nlm.nih.gov/). AutoDock Tools (version 1.5.6) was used for the activity pocket prediction of macromolecular proteins. Docking and free energy calculations of binding between AR and 3-demethylcolchicine were conducted using AutoDock Vina software. Finally, visualization was performed using Pymol (version 4.6.0) to effectively show the binding sites.

## 3. Results

### 3.1 Drug–compound–target network between Baihe and gastric cancer

The TCMSP database was searched for Baihe’s 84 active components, and 7 active compounds and 24 target genes were eliminated using the OB0.3 and DL0.18 filters **(Table 1)**. A total of 10,938 gastric cancer-related genes were obtained from GeneCards, OMIM, PharmGkb, TTD, and DrugBank databases. A total of 22 intersection genes were observed between Baihe targets and gastric cancer-related genes **(Figure 1)**. All of the seven active compounds, including isopimaric acid (MOL002039), stigmasterol (MOL000449), beta-sitosterol (MOL000358), 26-O-beta-D-glucopyranosyl-3beta (MOL000449), 3-demethylcolchicine (MOL009458), 26-O-β-D-glucopyranosyl-3β (MOL009465), and 26-O-β-D-glucopyranosyl-3β (MOL009471) were retained. A drug–compound–target network with seven compounds and 22 targets was generated using Cytoscape to illustrate the relationship of active compounds and disease targets intuitively **(Figure 2)**.

**Table 1.** Chemical components of Baihe

| Mol ID | Molecule Name | OB (%) | DL | Herb |
| --- | --- | --- | --- | --- |
| MOL009449 | 26-O-beta-D-Glucopyranosyl-3beta,26-dihydroxy-choleslen-16,22-dioxo-3-O-alpha-L-rhamnopyranosyl-(1-2)-beta-D-glucopyranoside_qt | 32.43 | 0.8 | Baihe |
| MOL009471 | 26-O-β-D-glucopyranosyl-3β,26-dihydroxy-cholestan-16,22-dioxo-3-O-α-L-rhamnopyranosyl-(1→2)-β-D-glucopyranoside_qt | 32.43 | 0.8 | Baihe |
| MOL009465 | 26-O-β-D-glucopyranosyl-3β,26-dihydroxy-5-cholesten-16,22-dioxo-3-O-α-L-rhamnopyranosyl-(1→2)-β-D-glucopyranoside_qt | 35.11 | 0.81 | Baihe |
| MOL002039 | Isopimaric acid | 36.2 | 0.28 | Baihe |
| MOL000358 | beta-sitosterol | 36.91 | 0.75 | Baihe |
| MOL009458 | 3-Demethylcolchicine | 39.34 | 0.57 | Baihe |
| MOL000449 | Stigmasterol | 43.83 | 0.76 | Baihe |

**Figure 1.**
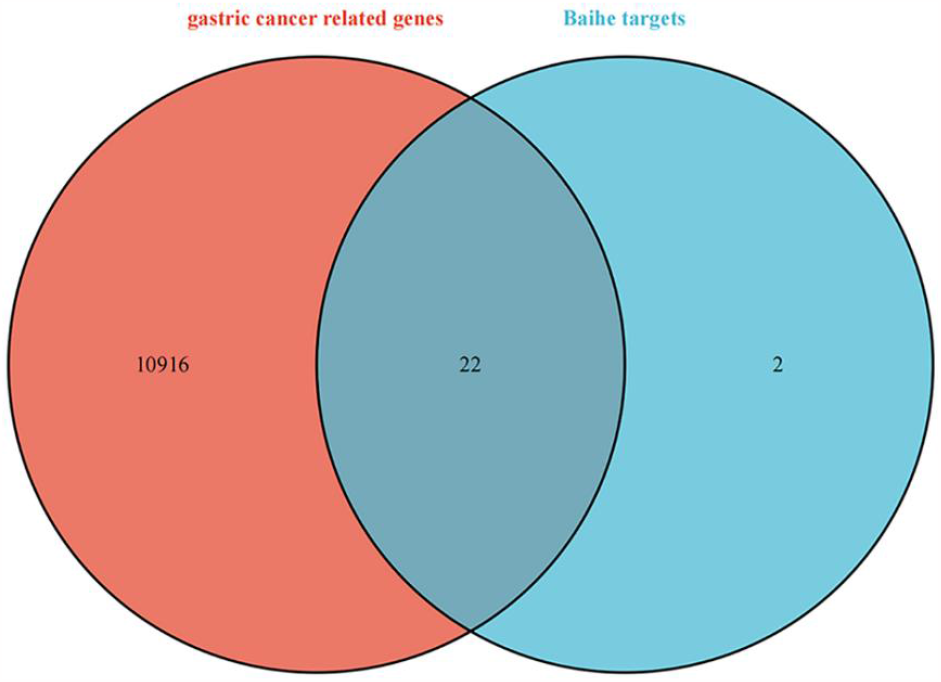
Venn diagram of the intersection between gastric-cancer-related genes and Baihe targets.

**Figure 2.**
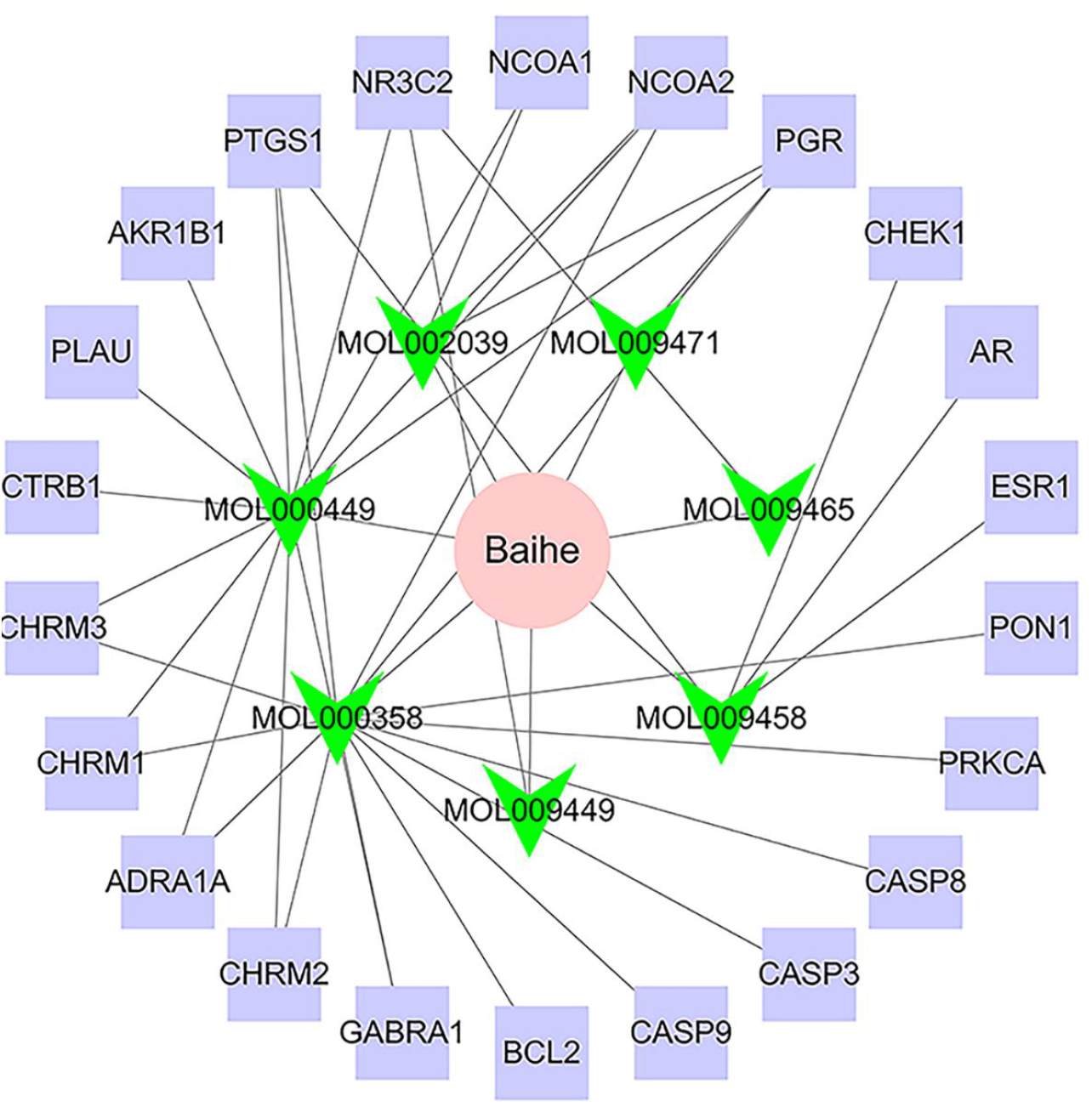
Active component-target-disease gene network.

The pink color represents Baihe, the green color represents the active component of Baihe, and the purple color represents the target of Baihe for gastric cancer treatment. The connected edges represent the relationship between the active components of the Baihe drug and the targets related to gastric cancer.

### 3.2 PPI network of 10 hub genes

The interaction of 22 overlapping proteins was verified by putting them into the STRING database (https://cn.string-db.org/), and 10 connected hub genes (AR BCL2 CASP3 ESR1 NCOA1 CASP8 NCOA2 CASP9 PGR NR3C2) satisfied the minimum required interaction score of 0.900 **(Figure 3)**. Three proteins (NCOA1, NCOA2, and CASP8) interacted with AR, indicating that AR proteins can interact with these three proteins and exert an effect on gastric cancer.

**Figure 3.**
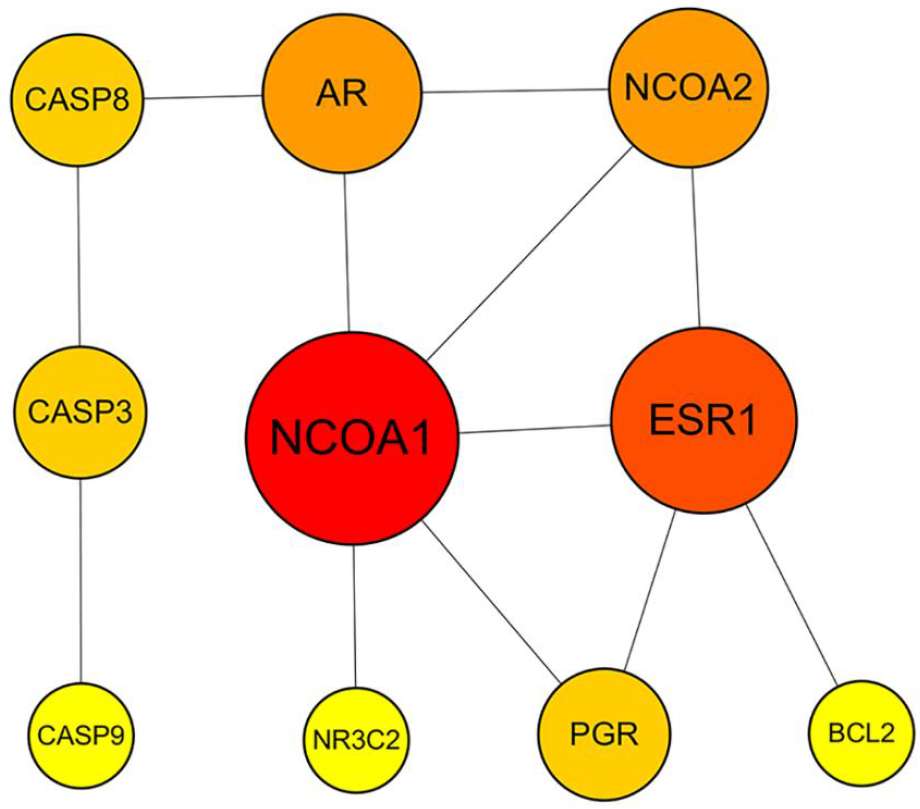
Protein–protein interaction network of Baihe.

The size and color of the circles represent the number of interacting proteins. The larger the circle, the more proteins interact with it; The closer the color is to red, the more proteins interact with it.

### 3.3 Functional enrichment and survival analyses of 10 hub genes

The pathways and biological function of 10 hub genes were determined by conducting KEGG and GO analyses to elucidate the mechanism of Baihe in the treatment of gastric cancer. The top seven entries in BP, MF, and CC in GO terms were retained **(Figure 4A-C)**, including “response to steroid hormone”, “cysteine−type endopeptidase activity involved in apoptotic signaling pathway”, “nuclear receptor activity”, and “transcription preinitiation complex”. The first 20 of 28 pathways were retained and shown in **Figure 5**, and main pathways included “p53 signaling pathway” and “apoptosis”.

**Figure 4.**
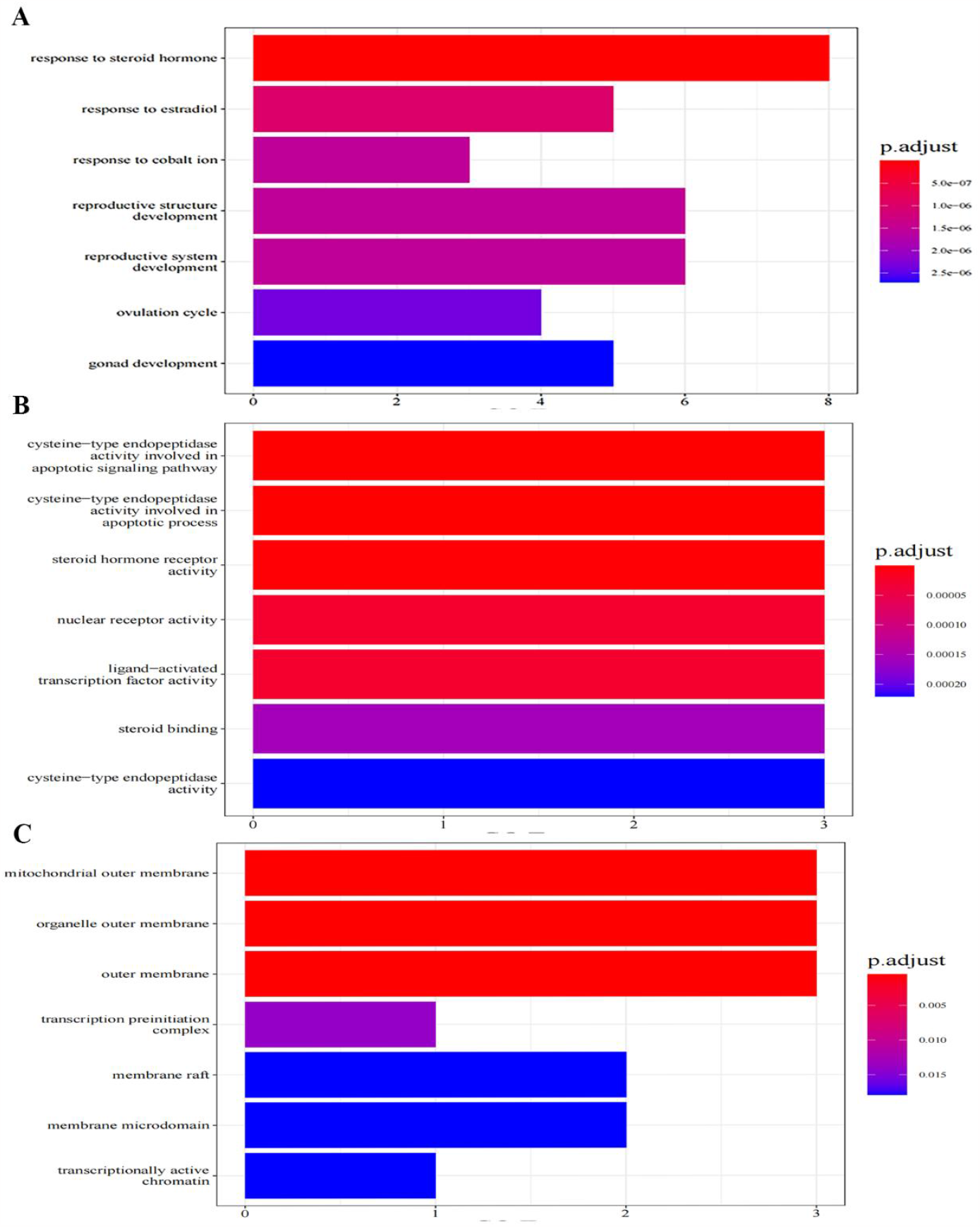
Gene Ontology (GO) enrichment analysis of the key genes of Baihe anti-gastric cancer. (A) GO biological process. (B) GO molecular function. (C) GO cellular components. The size of the dots represents the number of enriched genes, and the color represents the p adjustment.

**Figure 5.**
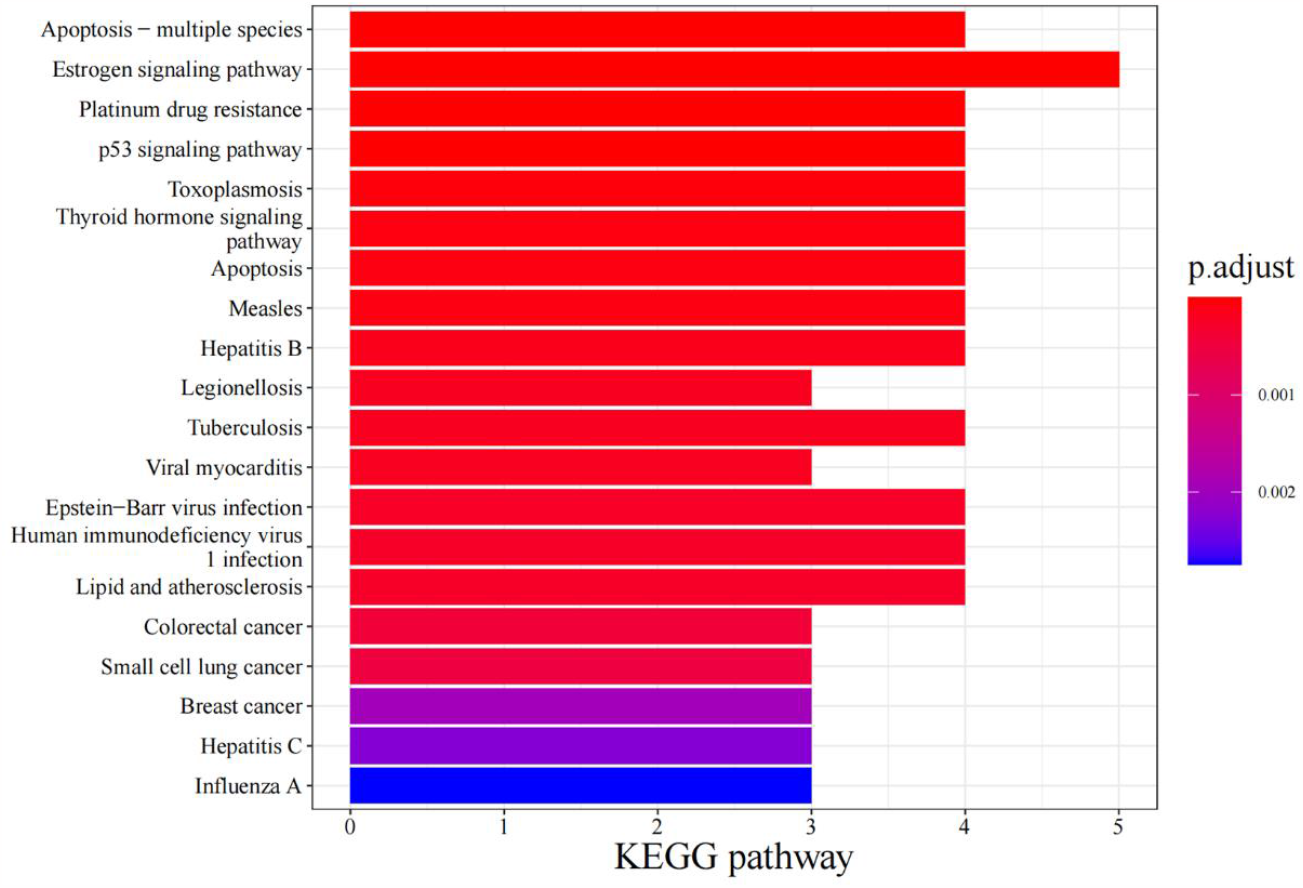
Kyoto Encyclopedia of Genes and Genomes pathway enrichment analysis of Baihe targets.

The length represents the enrichment gene number, while the color depth represents the extent of P adjust significance.

The prognostic effects of core genes on GC were investigated by conducting survival analysis on 10 gastric cancer hub genes in the Starbase database. Survival analysis results showed that the overall survival was significantly lower in patients with high AR expression than those with low expression (**Figure 6**, log-rank P< 0.05), while the differences in the results of other genes were not statistically significant (supplements). The literature review shows that key signaling molecules in gastric carcinogenesis were significantly regulated by AR activation (**Table 2**).

**Table 2.** The potential role of AR signals on the expression/activity of key molecules related to tumor growth in the gastric cancer

| Molecules | Main function | Mechanism | Reference |
| --- | --- | --- | --- |
| AKT<br>MMP9 | cell migration and invasion | AR-mediated up-regulation of MMP9 via activation of AKT | 43 |
| ZEB1 | cell metastasis and invasion | AR upregulated ZEB1 expression | 44 |
| EGFR | cell proliferation, migration and invasion | AR and EGFR may crosstalk | 45 |
| MYLK | cell migration | AR-v12 upregulated MYLK | 46 |
| LAMA4 | cell proliferation, migration and tumor initiation | AR activated LAMA4 expression | 47 |
| AURKA | cell cycle and several oncogenic pathways | AR and AURKA may crosstalk | 48 |
| CCRK | cell cycle | AR promote CCRK expression | 49 |
| Cyclin D1 | cell cycle | AR promote Cyclin D1 expression | 40 |
| P21 | cell cycle | AR promote P21 expression | 40 |
| P27 | cell cycle | AR inhibit P27 expression | 39 |
| P53 | cell cycle | AR and P53 may crosstalk | 41 |
| E-cadherin,<br>β-catenin | cell migration and invasion<br>cell cycle, proliferation and adhesion | AR inhibit E-cadherin expression<br>AR activated β-catenin expression | 37<br>37 |

**Figure 6.**
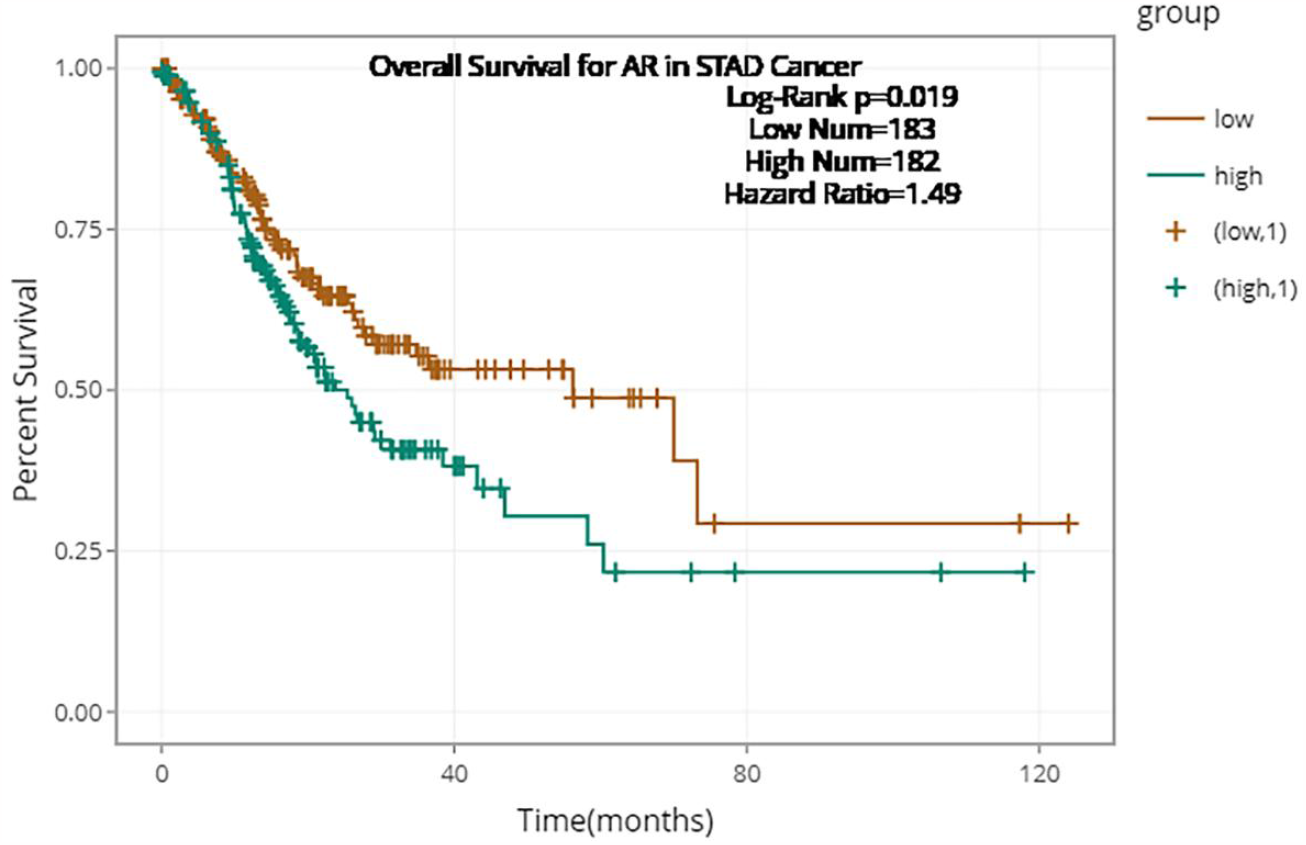
Kaplan-Meier curve of AR in GC cancer.

Survival analysis of AR in 365 patients with gastric cancer. The cut-off point for the high-risk group is the median. Brown represents 183 patients in the low-risk group, while green represents 182 patients in the high-risk group.

### 3.4 Molecular docking and visualization of AR and 3-demethylcolchicine

Among the seven active compounds and 22 targets, AR proteins and 3-demethylcolchicine were used for molecular docking and visualization because of the significant Kaplan-Meier survival analysis results of AR in gastric cancer. The mode of the binding sites of AR and 3-demethylcolchicine was predicted and showed by conducting molecular docking prediction and free energy calculation. Among the 20 predicted docking models, the top three minimum docking free energies were −6.7 kcal/mol (3 hydrogen bonds), −6.5 kcal/mol (2 hydrogen bonds), and −6.5 kcal/mol (2 hydrogen bonds), suggesting that the AR protein has a high affinity for the biological component 3-demethylcolchicine of Baihe. Accordingly, the top three docking models were visualized, and their docking sites are shown in **Figure 7**.

**Figure 7.**
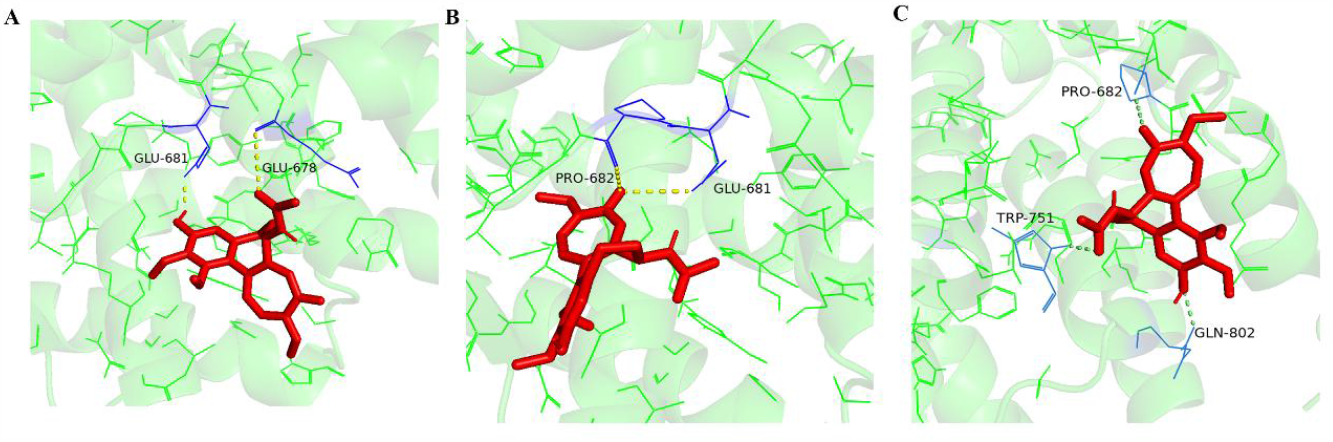
Docking model of compounds 3-demethylcolchicine with AR. The structure of the AR molecule is shown in green, the structure of 3-demethylcolchicine is shown in red, the hydrogen bonds that interact between the two are represented by the yellow dotted lines, and the amino acid residues linked by hydrogen bonds are shown in blue.

## 4. Discussion

As a result of its high mortality and morbidity, gastric cancer has become one of the most prevalent cancers. Clinically, patients with GC have poor prognosis because of the adverse drug reactions, drug resistance, and cancer recurrence^11^. Consequently, drugs that are safe and have few side effects need to be developed.

Many natural products derived from herbal medicine have gained considerable attention as antitumor drugs in recent years because of their high therapeutic value and low systemic toxicity ^12^. Baihe contains several bioactive compounds, including steroidal saponins, polysaccharides, flavonoids, phenols, glycosides, and steroidal alkaloids. These compounds have antitumor, antidepressant, antioxidant, anti-inflammatory, immunomodulatory, hypoglycemic, and antifungal effects^3, 13-15^. They include steroidal saponins, polysaccharides, and phenolics, which may have antitumor effects by interrupting the cycle, triggering apoptosis, and stimulating the immune system^14, 16^. However, the therapeutic effects and potential mechanisms of Baihe for GC are still unclear.

For the analysis and determination of the pharmacological mechanisms of Baihe for the treatment of GC, the combination of network pharmacology and molecular docking was used for the first time in this study. Seven significant Baihe-derived active compounds, including polysaccharide, isopimaric acid, beta-sitosterol, 3-demethylcolchicine, and stigmasterol, were used. Ten potential target genes such as AR, Bcl2, CASP3, ESR1, NCOA1, CASP8, NCOA2, CASP9, PGR, and NR3C2 were discovered. High AR expression was significantly linked to a poor prognosis for GC. Additionally, GO annotation and KEGG pathway enrichment analysis verified that these target genes are associated with responses to steroid hormones, cysteine-type endopeptidase activity involved in apoptotic signaling pathway, nuclear receptor activity, and transcription preinitiation complex, apoptotic, and p53 signaling pathways are associated, all of which are closely related to the treatment of GC. The main active compound of Baihe, 3-demethylcolchicine, has an affinity for the core target AR, according to the molecular docking results.

3-Demethylcolchicine, a colchicine metabolite, possesses a hydroxy-group on its carbon ring that could participate in radical scavenging and markedly inhibits the carrageenin edema^17^. Colchicine is an alkaloid that has been isolated from colchicum autumnale^18^. Colchicine can also be extracted from Baihe^19^. Colchicine has therapeutic effects in several diseases, including cancer ^20^, acute gout attacks, familial Mediterranean fever, leukoaraiosis, recurrent pericarditis, and primary biliary cirrhosis^21^. Moreover, colchicine has a narrow therapeutic window, since it is toxic, causes death at high doses, and exhibits significant toxicity to normal cell proliferation^22^. Colchicine is effective at doses of as low as 0.015 mg/kg, toxic at doses as high as 0.1 mg/kg, and lethal at doses as high as 0.8 mg/kg ^23, 24^. In terms of anticancer mechanisms, colchicine can prevent the proliferation of tumor cells by interacting with microtubules as a result of its anti-mitogenic activity^20^. Moreover, Lin et al. ^25^ found that the up-regulated DUSP1 gene may contribute to the anti-proliferative effect of colchicine on GC cells. Colchicine significantly inhibited tumor growth in nude mice by inducing apoptosis at doses of 0.05 and 0.1 mg/kg, and no visible toxicity was observed in liver and kidney tissues^26^. These results further illustrate the crucial role of colchicine for the treatment of GC.

The network pharmacology results demonstrated that the major 10 genes are the viable target genes of Baihe to modulate GC. The PPI network revealed close connections between these targets. The poor prognosis for GC was substantially related to the high expression of the target gene AR. The ligand-activated transcription factor AR, a member of the nuclear receptor superfamily and the steroid hormone receptor superfamily, is required for several cellular functions ^27^. When AR is expressed abnormally, it can act as an oncoprotein to cause the development and spread of many different types of cancer^28-32^. Hepatocarcinogenesis is fueled by the highly activated AR direct transcriptional target gene cell cycle-related kinase, which increases the *β-*catenin and T-cell factor signaling pathways^33^. The majority of prostate cancer is caused by aberrant AR activation, which is driven by mutations, amplifications, and shortened forms of AR^27^. The AR signaling axis is the target of the drugs abiraterone, enzalutamide, dalutamide, and apalutamide, which can enhance the prognosis of prostate cancer ^34^. In “in situ”, invasive, and metastatic breast cancer, AR is a common sex steroid receptor and is frequently overexpressed. AR-target drugs such as bicucultamide, abiraterone acetate, and enzalutamide have a significant role in the preclinical models of breast cancer^35^.

Aberrant AR expression is a major factor in the development and progression of gastric cancer^36^. According to epidemiological studies, men and women experience GC at a ratio of approximately 2:1 regardless of the etiology, which is related to how AR affects this disease ^37^. Shahrzad Soleymani Fard et al. found that 66.7% of patients with GC have high AR expression. High AR expression is associated to increased rates of lymphovascular invasion and lymph node metastasis, increased tumor mass, more distant AR expression, and a later TNM stage, which may significantly contribute to the poor prognosis of GC^38, 39^. By regulating the expression and protein activity of oncogene and antioncogene, AR can substantially affect the complex mechanism that causes and promotes the occurrence and advancement of GC^40-45^ (**Table 2**). For instance, AR promotes gastric carcinogenesis by interacting with genes related to cycles and EMT (P53, P21, P27, Cyclin D1, E-cadherin, and - catenin)^37, 46, 47^. Epidermal growth factor receptor (EGFR) belongs to the Erb B family of receptor tyrosine kinases (RTK) and is overexpressed in more than 30% of gastric adenocarcinomas. Considering the crosstalk between AR and EGFR pathways, a novel therapeutic strategy was proposed to inhibit the progression of GC by targeting both AR and subsequently EGFR pathways by using effective AR inhibitors such as enzalutamide^48^. These findings point to the critical role of AR in gastric carcinogenesis and the significance of targeting AR in the treatment of GC.

The potential biological functions of GC targets were identified by GO annotation and pathway enrichment analysis. GO enrichment of the main biological functions in response to steroid hormone, cysteine-type endopeptidase activity involved in apoptotic signaling pathway, nuclear receptor activity and transcription preinitiation complex. KEGG enrichment results showed that the p53 signaling pathway and apoptosis were the main signaling pathways in Baihe for GC treatment. Therefore, apoptosis and the P53 signaling pathway are key players in the development of GC. The P53 signaling pathway targets CDK, RPRM, p21, p16, TP53INP1, USF1/2, miR-17-5p, miR-20a, miR-181a, miR-449, and miR-650 to influence gastric cancer cell proliferation, differentiation, metastasis, cell cycle, apoptosis, immune response, and inflammation^49^.

Recent clinical studies have identified activated AR as a pathogenic feature of human malignant tumors, thus providing cancer cells with survival benefits. The activated AR can also be used for anti-cancer intervention. The molecular docking results show that 3-demethylcolchicine has a high affinity for AR, raising the possibility of a connection between Baihe bioactive substances and target genes.

The effect of 3-desmethylcolchicine on the AR signaling pathway has not been substantiated. Accordingly, a new therapeutic strategy was proposed for the treatment of GC that can target AR through 3-desmethylcolchicine, the bioactive compound in Baihe, for a highly effective anti-gastric cancer intervention.

## 5. Conclusion

In the present study, network pharmacology and molecular docking were used to clarify how Baihe regulates GC through various targets and pathways. The main target proteins of Baihe that are involved in GC have been demonstrated, resulting in the formation of a target protein network. Baihe mainly affected P53 and apoptotic signaling pathways while regulating AR key target proteins through the bioactive compound 3-demethylcolchicine.

Considering the complex components of Baihe, the data used in this article were collected using a partial database, and data need to be improved. The validity of the mechanism determined in the present study requires further research.

## Supporting information

supplement Fig1-9 Kaplan-Meier curve of target geneg in GC cancer.

## Supplementary Materials

The following supporting information can be downloaded at:

Figure S1: Kaplan-Meier curve of other target genes in GC cancer.

## Author contributions

Zi-Yi An and Wen-Hao Zhang performed the main analyses and draft the manuscript. Wei-Lin Jin designed the study. Xiao-Gang Hu helped with the introduction and discussion. All authors contributed to manuscript revision, read, and approved the submitted version.

## Institutional Review Board Statement

Not applicable.

## Informed Consent Statement

Not applicable.

## Data Availability Statement

The data presented in this study are available in this article and supplementary material.

## Acknowledgments

We thank the members of Jin Laboratory for their comments and discussions.

