## Supplementary figures and images for "Network pharmacological analysis for the identification of the molecular mechanism of *Lilium brownii* (Baihe) against gastric cancer: 3-Demethylcolchicine targeting androgen receptor"

### fig9.PGR.png

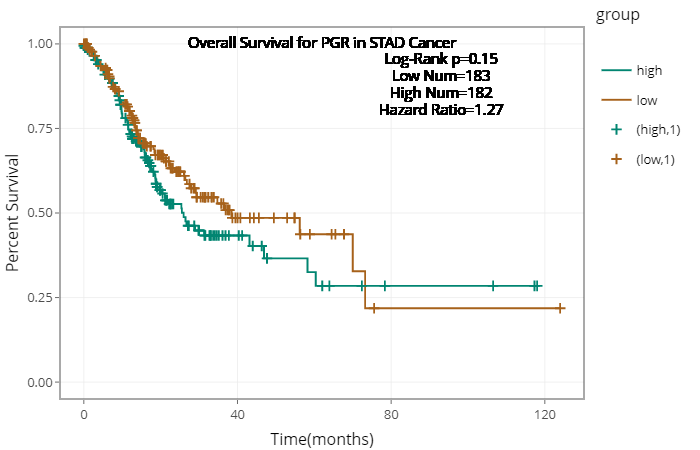

### fig 1.BCL2.png

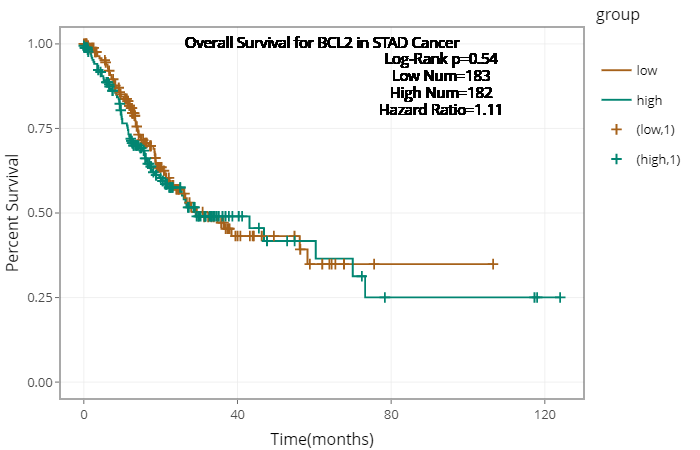

### fig 2.CASP3.png

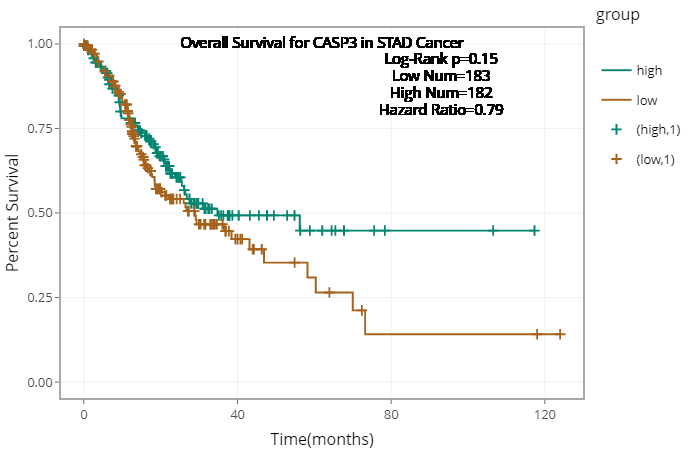

### fig 3.CASP8.png

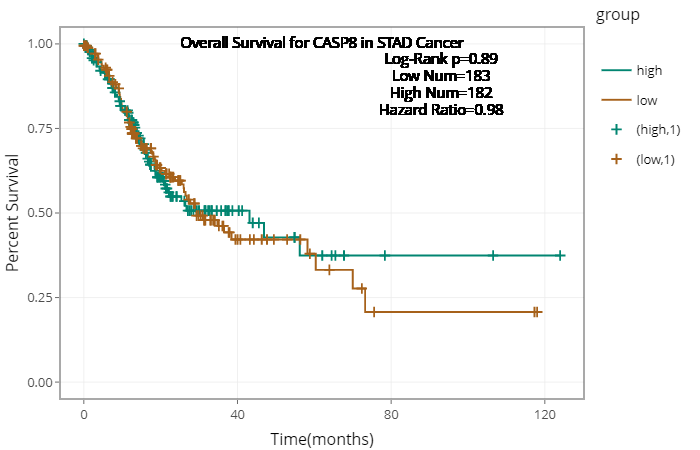

### fig 4.CASP9.png

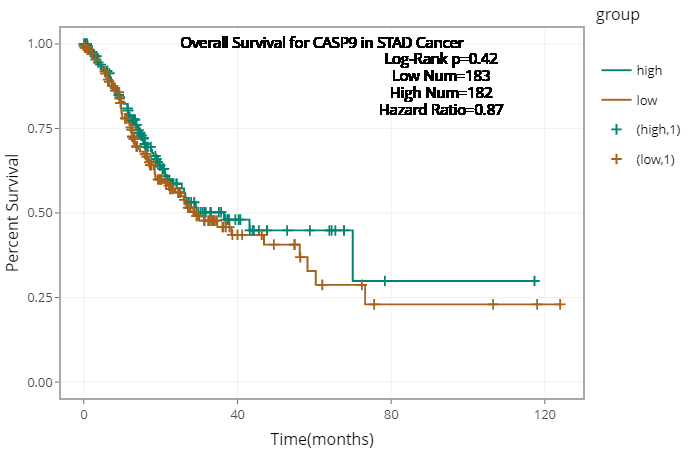

### fig 5.ESR1.png

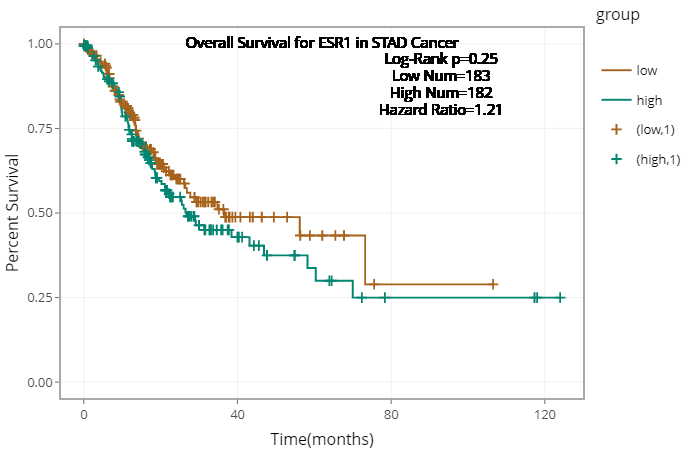

### fig 6.NCOA1.png

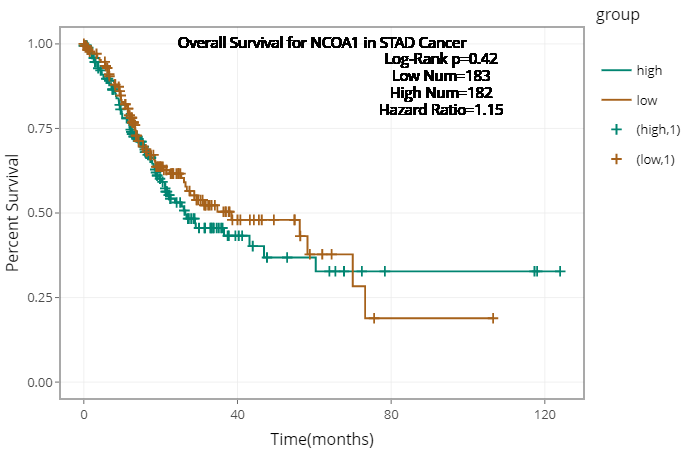

### fig 7.NCOA2.png

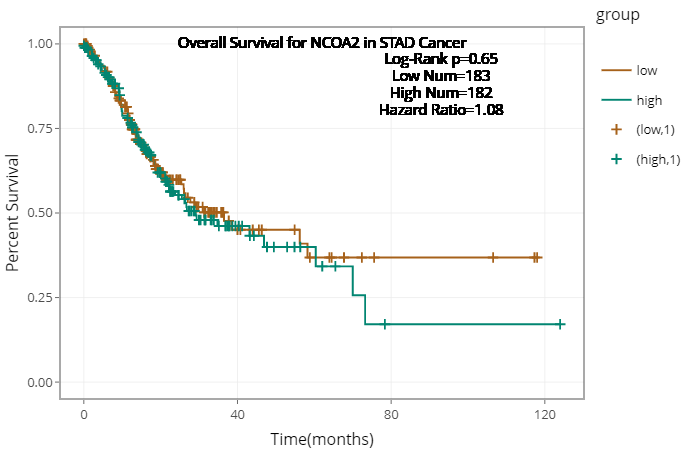

### fig 8.NR3C2.png

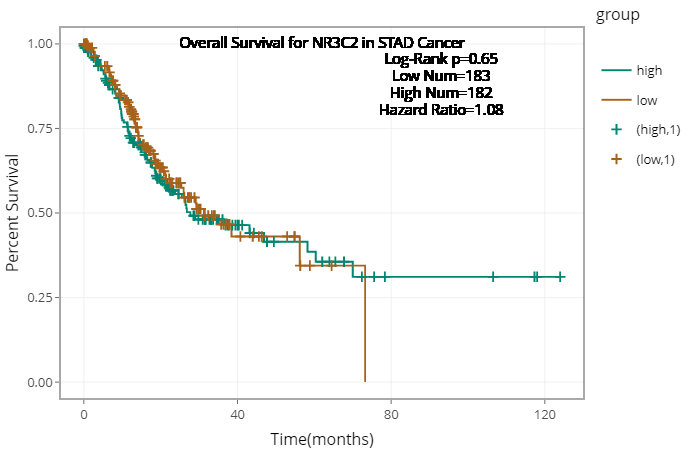
